# Selective activation of ganglion cells without axon bundles using epiretinal electrical stimulation

**DOI:** 10.1101/075283

**Authors:** Lauren E. Grosberg, Karthik Ganesan, Georges A. Goetz, Sasidhar Madugula, Nandita Bhaskar, Victoria Fan, Peter Li, Paweł Hottowy, Władysław Dabrowski, Alexander Sher, Alan M. Litke, Subhasish Mitra, E.J. Chichilnisky

## Abstract

Epiretinal prostheses for treating blindness activate axon bundles, causing large, arc-shaped visual percepts that limit the quality of artificial vision. Improving the function of epiretinal prostheses therefore requires understanding and avoiding axon bundle activation. This paper introduces a method to detect axon bundle activation based on its electrical signature, and uses the method to test whether epiretinal stimulation can directly elicit spikes in individual retinal ganglion cells without activating nearby axon bundles. Combined electrical stimulation and recording from isolated primate retina were performed using a custom multi-electrode system (512 electrodes, 10 µm diameter, 60 µm pitch). Axon bundle signals were identified by their bi-directional propagation, speed, and increasing amplitude as a function of stimulation current. The threshold for bundle activation varied across electrodes and retinas, and was in the same range as the threshold for activating retinal ganglion cells near their somas. In the peripheral retina, 45% of electrodes that activated individual ganglion cells (17% of all electrodes) did so without activating bundles. This permitted selective activation of 21% of recorded ganglion cells (7% of all ganglion cells) over the array. In the central retina, 75% of electrodes that activated individual ganglion cells (16% of all electrodes) did so without activating bundles. The ability to selectively activate a subset of retinal ganglion cells without axon bundles suggests a possible novel architecture for future epiretinal prostheses.

**New & Noteworthy:** Large-scale multi-electrode recording and stimulation were used to test how selectively retinal ganglion cells can be electrically activated without activating axon bundles. A novel method was developed to identify axon activation based on its unique electrical signature, and used to find that a subset of ganglion cells can be activated at single-cell, single-spike resolution without producing bundle activity, in peripheral and central retina. These findings have implications for the development of advanced retinal prostheses.

## 1 Introduction

Retinal prostheses are designed to restore partial visual function in patients blinded by photoreceptor degeneration. These devices operate by using electrode arrays to activate retinal neurons that have survived the degeneration process, causing retinal ganglion cells (RGCs) to transmit artificial visual signals to the brain. Clinically available prostheses are capable of generating visual percepts in patients using electrode arrays placed on the RGC side of the retina (epiretinal) (Humayun et al., 2012) or the photoreceptor side (subretinal) (Stingl et al., 2013b,a), with each approach exhibiting distinct advantages and disadvantages (Weiland et al., 2011; Goetz and Palanker, 2016). Subretinal implants can indirectly evoke spatially localized activity in RGCs by stimulating remaining inner retinal neurons, perhaps harnessing some of the visual processing capacity in the remnant circuitry (Lorach et al., 2015). However, stimulation of the different retinal cell types is indiscriminate and uncertain, and contrast sensitivity is low, perhaps as a consequence of indiscriminate stimulation (Goetz et al., 2015). Further, since the bipolar, horizontal, and amacrine cells that lie near subretinal implants are non-spiking, it is difficult to design a device that can record the elicited activity in order to fine-tune electrical activation and consequent visual signals transmitted to the brain. In contrast, direct activation of RGCs through epiretinal stimulation can elicit a wide variety of spike trains with high spatiotemporal precision and reproducibility (Jensen et al., 2003; Fried et al., 2006; Sekirnjak et al., 2008). With sufficiently high electrode density, selective activation of the appropriate RGCs at the correct times (Jepson et al., 2013, 2014a) could mimic the precise patterns of firing in 20 distinct RGC types that produce natural vision. Also, in principle, epiretinal implants could record elicited RGC spikes, in order to optimize stimulation patterns within the physical constraints of the interface.

However, a major challenge associated with epiretinal stimulation is activation of axon bundles in the nerve fiber layer, which lies between the electrodes and target RGCs. Large, arc-shape phosphenes that are elicited with the leading epiretinal implant (Argus II, Second Sight) almost certainly arise from axon bundle activation (Rizzo et al., 2003; Nanduri, 2011). One strategy to avoid bundle activation with epiretinal stimulation is to use long pulses or low-frequency sinusoidal stimulation to bypass RGCs and axons and instead stimulate the more distant retinal interneurons (Freeman et al., 2010; Boinagrov et al., 2014; Weitz et al., 2013, 2015). However, this approach eliminates the ability to producing precisely-controlled spike trains in multiple RGC types to mimic the natural output of the retina (Fried et al., 2006; Freeman et al., 2011). In principle, such precision is possible: epiretinal stimulation with high-density electrode arrays in peripheral retina has been shown to activate RGCs with single-cell, single-spike resolution (Sekirnjak et al., 2008; Jepson et al., 2013), emulating the native neural code of the retina in certain cases (Hottowy et al., 2012; Jepson et al., 2014a). However, for this approach to be viable in a prosthesis, it must be shown that high-precision stimulation can be achieved, to some degree, without axon activation. At present, it is not clear whether this is possible.

In this paper, we identify axon bundle activation in isolated primate retina on the basis of its characteristic electrical features recorded on a large-scale high-density multielectrode array. We then test whether high-precision somatic stimulation is possible in the absence of axon bundle activation, in peripheral and central retina. The results suggest that it may be possible to selectively activate a fraction of RGCs in parts of the central retina, with high precision and no bundle activation, using an epiretinal prosthesis. This raises the possibility of a novel and unique approach to optimizing the efficacy of artificial vision.

## 2 Materials and Methods

### Retinal preparation

Electrophysiology data were recorded from primate retinas isolated and mounted on an array of extracellular electrodes as described previously (Jepson et al., 2013). Eyes were obtained from terminally anesthetized rhesus macaque monkeys (*Macaca mulatta*, male and female, ages 4-20 years) used for experiments in other labs, in accordance with IACUC guidelines for the care and use of animals. After enucleation, the eyes were hemisected and the vitreous humor was removed. The hemisected eye cups containing the retinas were stored in oxygenated bicarbonate-buffered Ames’ solution (Sigma) during transport (up to 2 hours) back to the lab. Patches of intact retina 3 mm in diameter were placed in culture (Ames’ solution, gassed with 95% O_2_ and 5% CO_2_) with fluorescence conjugated peanut agglutinin or isolectin GS-IB4 (AlexaFluor 568, cat. nos. L32458 or I21412, Life Technologies) at 4.14 µg/mL (36.5pM) for 5-10 hours for vasculature labeling before proceeding with the dissection. Then, the retina was isolated from the pigment epithelium under infrared illumination and held RGC-side down on a custom multi-electrode array (see below). Throughout the experiments, retinas were superfused with Ames’ solution at 34°C.

### Electrophysiological recording and stimulation

A custom 512-electrode stimulation and recording system (Hottowy et al., 2008, 2012) was used to apply electrical stimuli and record spikes from RGCs. The electrode array consisted of 512 electrodes in a 16x32 isosceles triangular lattice arrangement, with 60 µm spacing between electrodes within rows, and between rows (Litke et al., 2004). Electrodes were 10 µm in diameter and electroplated with platinum. For recording, raw voltage signals from the electrodes were amplified, filtered (43-5000 Hz), and multiplexed with custom circuitry. These voltage signals were sampled with commercial data acquisition (DAQ) hardware (National Instruments) at 20 kHz per channel. For stimulation, custom hardware (Hottowy et al., 2012) was controlled by commercial multifunction DAQ cards (National Instruments). Charge-balanced triphasic current pulses with relative amplitudes of 2:-3:1 and phase widths of 50 µs (total duration 150 µs) were delivered through one electrode at a time. Reported current amplitudes correspond to the magnitude of the second, cathodal, phase of the pulse. This pulse shape was chosen to reduce stimulation artifact in the recordings. Custom artifact reduction circuitry disconnected the recording amplifiers during stimulation, reducing stimulation artifact and making it possible to identify elicited spikes on the stimulating electrode and nearby electrodes (Hottowy et al., 2012; Jepson et al., 2013). For recording and stimulation, a platinum ground wire circling the recording chamber served as a distant ground.

### Electrical images and cell type classification

Recordings obtained with visual stimulation were analyzed to identify spike waveforms of distinct RGCs in the absence of electrical stimulation artifact, using spike sorting methods described previously (Litke et al., 2004; Field et al., 2007), which identified spike times of identified RGCs based on the relatively large, stereotyped spikes detected near the soma. Then, the complete spatio-temporal signature of the spikes from each cell over all electrodes, or electrical image, was computed, by averaging the voltage waveforms on all electrodes at and near the times of its recorded spikes (Litke et al., 2004) (Fig. 2). The electrical image of each cell provided a template of its spike waveform, which was used to identify the cells producing spikes in response to electrical stimulation (Fig. 1E).

**Figure 1:**
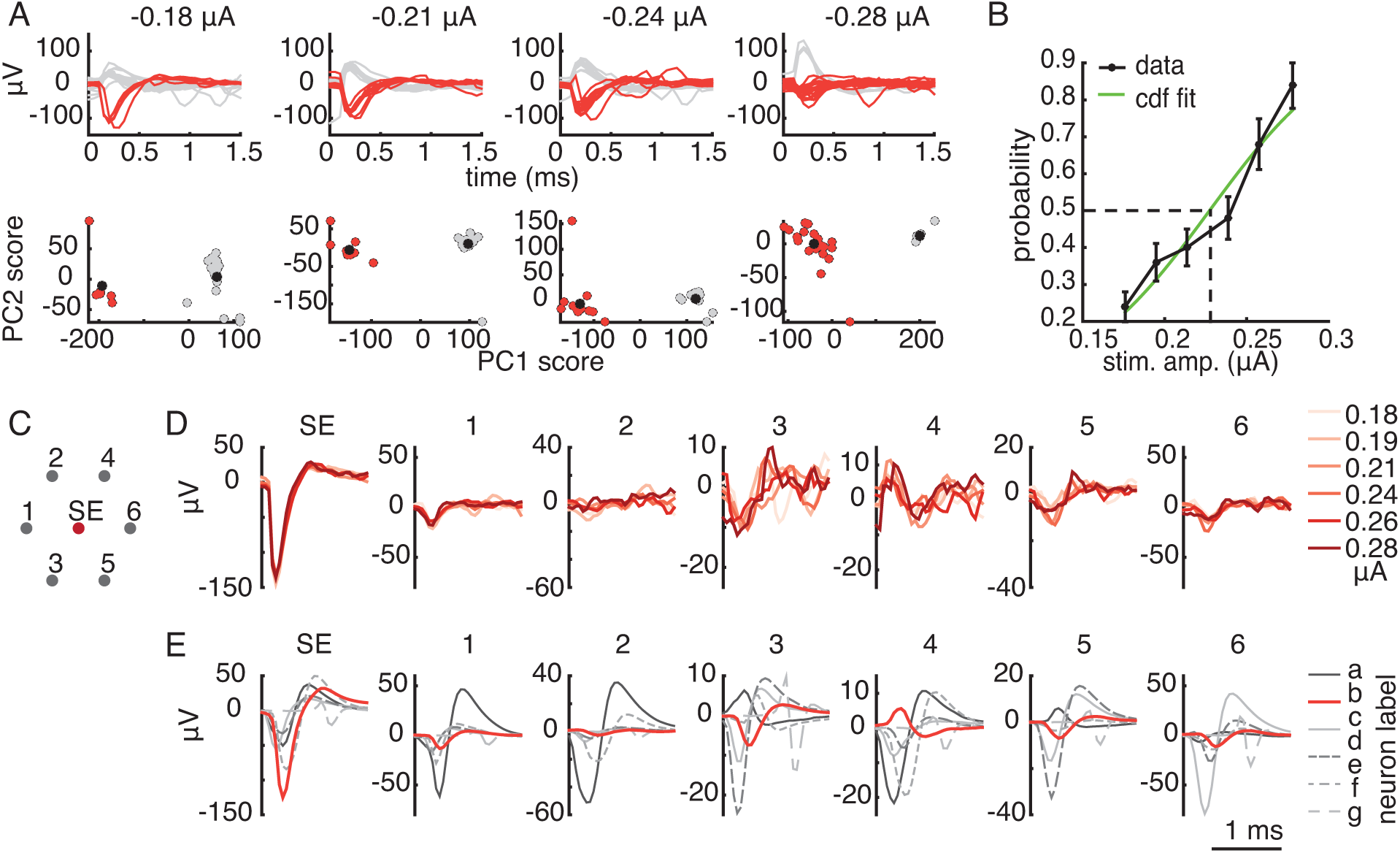
Semi-automated method for detecting somatic activation. A. Top row: mean-subtracted waveforms recorded on the stimulating electrode immediately following electrical stimulation, at four stimulation amplitudes. Bottom row: at the same amplitudes, the coefficients for each trial corresponding to the first 2 principal components of the recorded waveforms form distinct clusters. Estimated cluster centers are indicated by black circles. Red (gray) waveforms and points indicate trials that were identified automatically as containing (not containing) spikes. B. A cumulative Gaussian function was fitted to the probability of activation across trials computed from the data in A. Error bars represent ± one standard deviation of 100 probabilities computed after resampling the trials with replacement 100 times. The activation threshold (0.23 µA) was defined as the stimulation amplitude that produced 50% activation probability according to the fitted function. C. Electrode configuration for the waveform comparison. Recordings from the stimulating electrode (SE) and the surrounding electrodes were inspected. Numbers correspond to the waveform plots in D-E. D. The consistency of the waveform shape at consecutive stimulation amplitudes was verified to ensure that the elicited waveform was produced by a single cell. For each stimulation amplitude (indicated by the shade of red in the plot), the mean of the non-spiking trials was subtracted from the mean of the spiking trials on the stimulating and surrounding electrodes to extract the electrically elicited spikes waveform. E. Spike templates (electrical images) obtained with visual stimulation reveal 7 RGCs recorded at the stimulating electrode, labeled by their unique neuron labels a-g. The template waveform (b) that matched the electrically elicited waveforms shown in D is highlighted in red.

**Figure 2:**
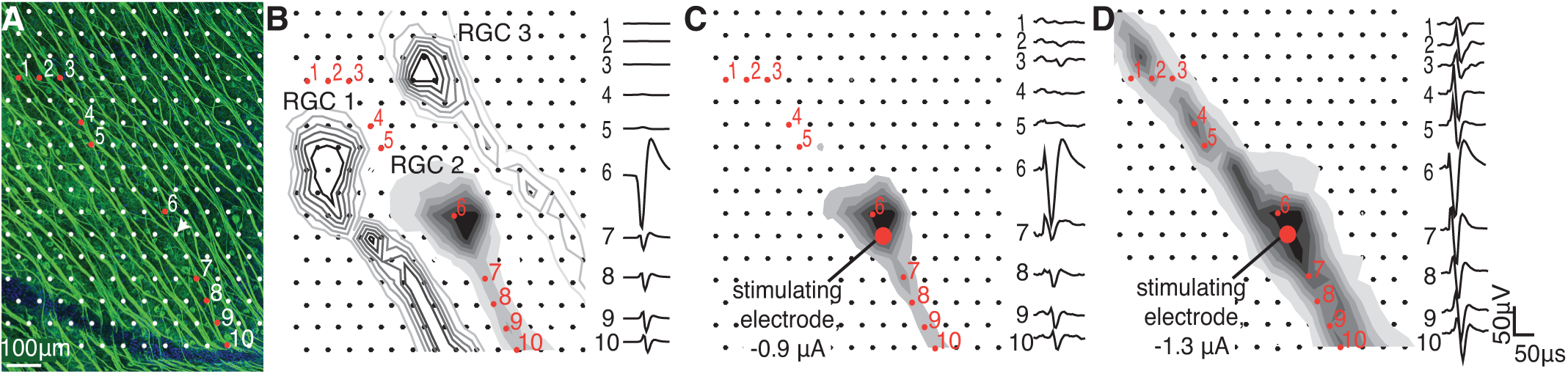
Bidirectional propagation of electrically evoked responses. A. Fluorescence image shows the density and arrangement of RGC axon bundles with respect to the electrode array. Arrow indicates the stimulating electrode in panels C-D. B. Electrical images (EIs) from three RGCs in a single retina obtained with visual stimulation (no electrical stimulation). All of the axons run in the same direction toward the optic disc. Waveforms on the electrodes indicated with numbers are associated with the shaded RGC. C-D. Unidirectional (C) and and bi-directional (D) signal propagation after electrical stimulation at the electrode shown by a red dot. Note similarity of unidirectional (C) image to the EI from the shaded cell shown in B. The amplitude of the waveform shown for electrode 6 in all panels was reduced by a factor of 2 relative to the scale bar.

Distinct RGC types were identified by their distinct responses to white noise visual stimuli. Briefly, a dynamic random checkerboard stimulus was presented, and the average stimulus that preceded a spike in each RGC was computed, producing the spike-triggered average (STA) stimulus (Chichilnisky, 2001). The STA summarizes the spatial, temporal and chromatic properties of light responses. Features of the STA were used to segregate functionally distinct RGC classes. Spatial receptive fields (RFs) for each cell type (Fig. 8) were obtained from fits to the STA (Chichilnisky and Kalmar, 2002). For each identified RGC type, the receptive fields formed regular mosaics covering the region of retina recorded (Devries and Baylor, 1997; Field et al., 2007), confirming its correspondence to a morphologically distinct RGC type (Wassle et al., 1981; Dacey, 1993), and in some cases revealing complete recordings from the population. The density and light responses of the four most frequently recorded RGC types uniquely identified them as ON and OFF midget, and ON and OFF parasol, which collectively account for 68% of RGCs in primates (Dacey, 2004). Other RGC types were encountered but not identified. The regular mosaic structure of RGC receptive fields of each type (Devries and Baylor, 1997; Chichilnisky and Kalmar, 2002; Gauthier et al., 2009) was used to estimate the total number of cells expected to be present over the array (Table 1). Because OFF midget cell mosaics were sparse in the recordings presented, the number of OFF midget cells was estimated as follows: *m*_(−)_ = *m*_(+)_*p*_(−)_*/p*_(+)_, where *m*_(−)_, *m*_(+)_, *p*_(−)_, and *p*_(+)_ represent the number of OFF midget, ON midget, OFF parasol and ON parasol cells over the array, respectively. The total number of RGCs expected to be present over the array was estimated as (*m*_(−)_ + *m*_(+)_ + *p*_(−)_ + *p*_(+))_/0.68 (Dacey, 2004).

**Table 1:** Summary of selective activation results in three peripheral recordings. Percentages in parentheses obtained by dividing the data in each entry by the entry immediately above. The category labeled “other” includes OFF midget cells (see (Gauthier et al., 2009)), small bistratified cells (see (Field et al., 2007)), and cell types for which the anatomical identity is unknown.

| eccentricity | counts | ON parasol | OFF parasol | ON midget | other | total |
| --- | --- | --- | --- | --- | --- | --- |
| 48.2° | # total RGCs present | 119 | 177 | 298 | 931 | 1525 |
|  | # recorded RGCs | 116 (0.97) | 150 (0.85) | 202 (0.68) | 68 (0.07) | 538 (0.35) |
|  | # single RGC activation | 78 (0.67) | 69 (0.46) | 43 (0.21) | 29 (0.43) | 219 (0.41) |
|  | # selective activation | 37 (0.47) | 27 (0.39) | 29 (0.67) | 19 (0.66) | 112 (0.51) |
| 58.1° | # total RGCs present | 55 | 72 | 287 | 747 | 1161 |
|  | # recorded RGCs | 48 (0.87) | 34 (0.47) | 211 (0.74) | 80 (0.11) | 373 (0.32) |
|  | # single RGC activation | 41 (0.85) | 12 (0.35) | 36 (0.17) | 37 (0.46) | 126 (0.34) |
|  | # selective activation | 25 (0.61) | 3 (0.25) | 23 (0.64) | 20 (0.54) | 71 (0.56) |
| 58.1° | # total RGCs present | 51 | 64 | 279 | 700 | 1094 |
|  | # recorded RGCs | 42 (0.82) | 29 (0.45) | 205 (0.73) | 56 (0.08) | 332 (0.30) |
|  | # single RGC activation | 40 (0.95) | 23 (0.79) | 67 (0.33) | 28 (0.50) | 158 (0.48) |
|  | # selective activation | 22 (0.55) | 8 (0.35) | 39 (0.58) | 6 (0.21) | 75 (0.47) |
| all | # total RGCs present | 225 | 313 | 864 | 2378 | 3780 |
|  | # recorded RGCs | 206 (0.92) | 213 (0.68) | 618 (0.72) | 204 (0.09) | 1243 (0.33) |
|  | # single RGC activation | 159 (0.77) | 104 (0.49) | 146 (0.24) | 94 (0.46) | 503 (0.40) |
|  | # selective activation | 84 (0.53) | 38 (0.37) | 91 (0.62) | 45 (0.48) | 258 (0.51) |

### Identification of axon bundle activation

To analyze electrically evoked activity over the entire array, the voltages recorded on all electrodes immediately following stimulation were obtained, with 15-25 trials (repeats) of each electrical stimulus condition. Mean voltage waveforms recorded with the lowest stimulation current amplitude for a particular electrode were subtracted from data recorded with all higher stimulation amplitudes at that electrode, to reduce (but not eliminate) the electrical stimulus artifact. Then, in order to identify axon bundle activation, either human inspection or an automated algorithm was used.

For human inspection, the observer viewed movies of the recorded activity following stimulation and identified the lowest current amplitude at which bidirectional propagation was visible. Analyzing the results from six preparations required viewing *>*15,000 movies of activity for all different electrodes and stimulus amplitudes.

For automated bundle detection, the responses to electrical stimulation were mapped to a collection of weighted graphs, and graph partitioning and graph traversal algorithms were applied to identify bundle activity. The focus was on two characteristic features of axon bundle signals: bidirectional propagation, and growth of signal amplitude with stimulation current. The algorithm, and an evaluation of its sensitivity with synthetic data, are described in detail in the Appendix.

### Identification of somatic activation

Voltage traces recorded on the stimulating electrode immediately following stimulation were analyzed to find the lowest current amplitude that produced reproducible somatic activation. We use this term to refer to activation by an electrode that records a somatic spike from the cell, as revealed by the characteristic large voltage deflection that begins with a dominantly negative component. Note that it is possible and indeed likely that the site of activation is actually the axon initial segment (Sekirnjak et al., 2008; Fried et al., 2009). To obtain the somatic activation thresholds in a semi-automated way, two features of electrical stimulation data were utilized: elicited spikes are stochastic for stimulation current levels near activation threshold for a given RGC, and the timing of elicited activation is consistent across trials (Sekirnjak et al., 2008; Jepson et al., 2014b). Therefore, recorded voltage traces following stimulation were divided into distinct groups when there was partial RGC activation, as shown in Fig. 1A (top row). Principal component analysis (PCA) across trials was used at each stimulation amplitude, and clusters in principal component (PC) space were used to identify the amplitudes in which trials fell into distinct groups (Fig. 1A, bottom row). A fuzzy c-means clustering algorithm (MATLAB and Fuzzy Logic Toolbox Release 2014b, The MathWorks, Inc.) was used to find two clusters in the first two PCs of the data. Each PC data point was assigned a membership grade (between 0 and 1) for each cluster based on distance to the cluster centers. A threshold on membership grades for all points in a cluster was used to identify which stimulation amplitudes produced two distinct clusters, indicating the presence of spikes on some trials but not others. For a cluster to be considered “distinct”, all members were required to have membership grades of 0.8 or higher, with 2 outliers allowed. These parameters were chosen so that the analysis erred towards positively identifying clusters, minimizing false negatives. Clustering was confirmed by human inspection to remove false positives, and a cumulative Gaussian function was fit to the response probabilities to determine the 50% activation threshold (Fig. 1B).

After identifying RGC spikes for current levels near the threshold of activation, the remaining analysis steps were to determine whether the elicited waveforms over the range of stimulation amplitudes were consistent with spikes from a single cell, and if so, to identify the cell type of the activated cell. These steps were performed by human inspection. First, the mean of the non-spiking trial responses was subtracted from the mean of the spiking trial responses for the stimulating electrode and surrounding electrodes. The resulting residual waveform was checked for consistency across stimulation amplitudes, which would be expected from the activation of a single cell (Fig. 1D). Responses producing multiple distinct residual waveforms were counted as failures of selective activation (Figs. 7–9). Next, these residual waveforms (Fig. 1D) were compared to the template waveforms (electrical images) of cells recorded with visual stimulation (Fig. 1E), to identify the cell and cell type of origin. Responses from cells not identified this way were counted as “other” cell types (Fig. 8).

### Immunohistochemistry and imaging

Following stimulation and recording, wide-field fluorescence imaging was used to visualize axon bundles in the recorded piece of retina. Immunolabeling was performed as described previously (Li et al., 2015). Tissue was fixed with 4% paraformaldehyde in phosphate buffered saline (PBS, 10 mM) for 45 minutes at room temperature and then washed in PBS 3x10 minutes and left in PBS at 4deg C for 6-48 hours. Fixed tissue was incubated for 2 hours in blocking solution at room temperature, and then incubated in blocking solution with a 1:200 dilution of primary antibody (rabbit monoclonal anti-βIII-tubulin, Abcam cat. no. ab52623 or Covance cat. no. MRB-435P) for 2-3 days at 4deg C on a shaker. The blocking solution consisted of 10% normal donkey serum and 0.25% Triton X-100 in PBS. In some cases, 0.05% sodium azide was added as a preservative. Following incubation with primary antibody, the tissue was washed in PBS 3x10 minutes. Tissue was then incubated in blocking buffer with fluorescence-conjugated (either AlexaFluor 488 or Cy3) secondary antibodies (goat anti-rabbit IgG H&L, either Abcam or Jackson ImmunoResearch) at a 1:200 dilution for 3-4 hours at room temperature on a shaker, washed in PBS 3x5 minutes, counterstained with DAPI 10 minutes, and mounted on slides in ProLong Gold antifade medium (Life Technologies). Coregistration of the electrode array to the fluorescence image (Fig. 2A) was performed using the labeled vasculature before and after fixation as described in (Li et al., 2015).

## 3 Results

A high-density 512-electrode system (Hottowy et al., 2012) was used to electrically stimulate and record neural activity in isolated macaque retina. To determine the origin of the recorded activity, the response to electrical stimulation was compared to the electrical signatures of individual RGCs and axons obtained with visual stimulation, and to fluorescence images of axon bundles, in the same preparation. The thresholds for somatic and axonal activation were then examined to determine how selectively RGCs could be activated without axonal activation, and the implications for design of future epiretinal prostheses.

### Bundle activation can be identified by its electrical signature

To identify the electrical signatures of RGCs and axons overlying the electrode array, the electrical image of every RGC recorded during visual stimulation with spatiotemporal white noise was calculated (see Methods). The electrical image of a cell is the average spatiotemporal voltage pattern produced across the array during a spike (Petrusca et al., 2007; Greschner et al., 2014). All RGCs in each preparation had electrical images comprised of signals that apparently initiated near their somas and propagated along their axons en route to the optic disc (Fig. 2B). As expected in peripheral recordings, the axons in the electrical images were approximately parallel to one another. An image of axon bundles obtained with antibodies to tubulin after recording (see Methods) revealed that the orientation of axon bundles corresponded closely to the orientation of the axons inferred from the electrical images (Fig. 2A).

Responses to electrical stimulation were then compared to these electrical images, in some cases revealing the activation of an individual RGC. This was observed by examining the mean voltage deflection recorded on each electrode after passing current through a selected electrode, after subtraction of the artifact at the lowest stimulation amplitude (see Methods). In general, a large signal, composed of physiological signal and stimulus artifact, was observed near the stimulating electrode immediately after the stimulus (Fig. 2C,D). A voltage deflection then propagated away from the stimulating electrode. In some cases, the starting point and unidirectional propagation of the elicited activity closely matched the electrical image of one of the RGCs obtained during visual stimulation, indicating that this specific cell was activated in isolation (Fig. 2C). In other cases, the recorded signal originated at the stimulating electrode and propagated bidirectionally along the direction of the axon bundles (confirmed by imaging, Fig. 2A), indicating that passing axons of one or more RGCs had been activated by the electrical stimulus (Fig. 2D). In both cases, the speed of propagation was 1.1 ± 0.39 m/s (measured on 1245 electrodes in 3 retinas), consistent with action potential propagation speeds recorded from single axons measured using the same experimental methods (Li et al., 2015). However, in the case of bidirectional propagation, the spatial pattern of the recorded signal did not match the pattern of any single electrical image obtained in the preparation (e.g., Fig. 2D). This mismatch suggests that the signal was comprised of axons of multiple RGCs, and/or axons not identified during visual stimulation.

In the case of bidirectional propagation, several additional aspects of the recorded voltage signal provided information about the cells and axons activated. The average amplitude of the recorded waveforms increased with stimulation current, and the responses in different trials with the same stimulation current had similar amplitudes (Fig. 3C). These observations are consistent with the progressive recruitment of multiple axons, and inconsistent with the all-or-none signals that would be produced by a spike in a single neuron. Furthermore, stepwise increases in response amplitude (Fig. 3C) suggested that different axons were recruited at different stimulation current levels. Cross-sectional voltage profiles (orthogonal to the direction of axon propagation) (Fig. 3A) were wider at higher stimulation current levels (Fig. 3B), suggesting the recruitment of axons laterally displaced from the stimulating electrode and perhaps comprising different bundles. These observations were consistently observed at the large majority of stimulating electrodes tested in eight preparations.

**Figure 3:**
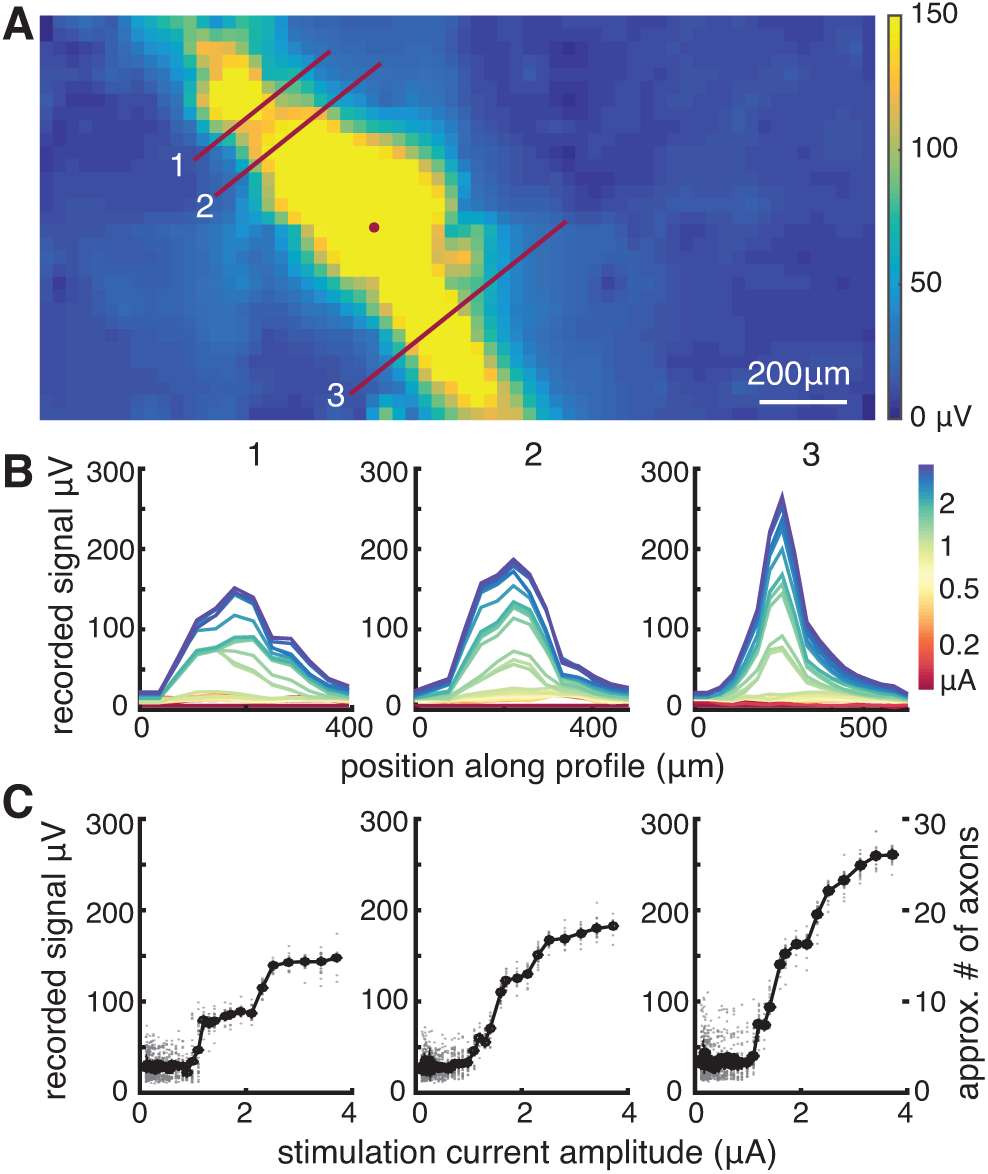
Axon recruitment in bidirectional axon bundle signal. A. 2D interpolation of the maximum amplitudes recorded in the 5 ms after stimulation at the electrode indicated by the circle (mean of 25 trials). Lines indicate cross sections over which amplitudes are shown in B-C. The ‘downstream’ signal direction toward the optic nerve is down-right, the ‘upstream’ direction is up-left. B. Voltage recordings along the profiles shown in A. Numbers correspond to the profile line. Colors indicate stimulation amplitude for each voltage profile shown. C. The maximum amplitude of the profiles (averaged across trials) shown in B increase progressively with stimulation current. Data from individual trials is shown in gray. The approximate number of axons activated, based on the amplitude of a typical axon in this retina, is shown for comparison (far right).

The number of axons in the evoked signal was estimated by comparing the response obtained at high stimulus amplitudes to the mean single-axon response amplitude, obtained from electrical images of RGCs identified with visual stimulation in the same preparation. For example, in one case (Fig. 3C, right panel) the peak voltage deflection observed in response to the maximum amplitude stimulation (3.7 µA), which produced a saturating response, was 27-fold larger than the mean voltage deflection associated with individual identified axons in the same preparation. Over all electrodes along the activated path, shown in yellow in Fig. 3A, the mean recorded signal was 13-fold larger than the average single axon signal amplitude. In the same preparation, the mean of the signals recorded along the activated paths produced after stimulation at each of the 512 electrodes was 12-fold (± 5-fold, 1 SD) larger than a single axon signal. Note that these estimates of the number of axons could be biased by the spatial arrangement of axons stimulated and recorded, and the selection of cells for which electrical images were obtained and used for single-axon amplitude estimates. With this caveat, the results suggest that tens of axons from several nearby bundles were activated at the maximum stimulus amplitude, and that the stepwise activation of axons probably reflected groups of axons, perhaps comprising different bundles, being recruited at specific current levels.

On the basis of these observations, in what follows the evoked bidirectional signal will be referred to as axon bundle activation, although it is likely that in some cases only one axon was activated at low stimulus current levels (see Fig. 6).

### Axon bundle activation thresholds vary over electrodes and preparations

The form of the current-response curve permitted the identification of a threshold for bundle activation. Specifically, in most cases the recorded signal amplitude exhibited a discrete change in slope at a particular stimulation current level (e.g., 1.2 µA in Fig. 3B-C). The threshold amplitude for the activation of bundles was identified for all electrodes (excluding array borders) in each preparation by examination of the voltage recordings after a stimulation pulse, focusing on the presence of bidirectional propagation (see Methods). The threshold for activation was defined as the midpoint between successive stimulation amplitudes at which bidirectional activity was and was not observed. Bundle threshold typically varied several-fold across electrodes (Fig. 4). For six preparations from five retinas, the mean axon bundle activation threshold was 1.34 ± 0.58 µA (mean ± SD). This value is comparable to the thresholds for somatic stimulation of individual RGCs (see below) (Jepson et al., 2013).

**Figure 4:**
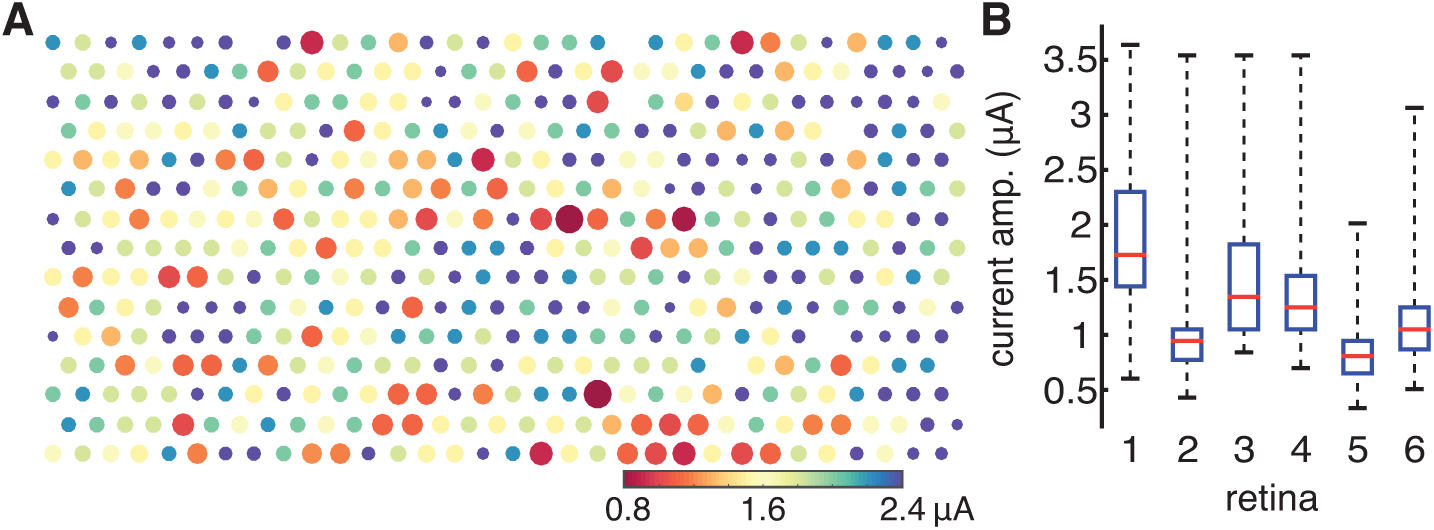
Axon bundle activation threshold map exhibits spatial variability. A. In a single recording, bundle threshold for each electrode is indicated by the color of the dot representing that electrode and the inverse of the dot size. B. For six preparations from five retinas (3 and 4 represent different preparations from the same retina), box-and-whisker plots summarize the medians, quartile values, and ranges of the thresholds for all electrodes in the recording. Data from retina 1 is shown in A.

Different recordings from peripheral retina exhibited substantially different axon bundle thresholds. For each of the six preparations described above, the observed thresholds were (in µA): 1.87 ± 0.58, 0.96 ± 0.35, 1.48 ± 0.51, 1.41 ± 0.50, 0.84 ± 0.26, and 1.14 ± 0.41 (mean ± SD across electrodes, in each preparation). The differences between preparations were statistically reliable. This was determined by pooling the thresholds obtained in all six recordings, resampling with replacement from this pool to create six distinct groups, each containing the same number of observations as one of the preparations, and computing the RMS difference between the means of the groups. The observed value of 2.07 in the real data was well outside the range of the resampled data, 0.16 ± 0.11 (mean ± 2SD), indicating that the thresholds from different preparations were not drawn from a single population. Diverse factors could contribute to this variation, including spatial variations in the axon layer thickness and pressing of the retina against the electrode array.

An automated algorithm for bundle activation detection (see Methods) provided estimates of activation threshold similar to those identified by manual analysis. Comparison of the manual and automated analysis revealed a 0.93 correlation between the methods over 1885 electrodes examined in five preparations from four retinas (Fig. 5A). 62% of the thresholds identified were identical, and 88% of the thresholds agreed within ±1 experimental stimulation amplitude step (each step corresponded to a 10% increment in stimulation current, Fig. 5B). For this analysis, stimulation electrodes at the border of the array were ignored, because of the difficulty of detecting bidirectional propagation at the border in manual analysis. Importantly, the algorithm captured the growth of the recorded signal with stimulation amplitude (as shown in Fig. 3C), and used it to help identify bundle activation, instead of relying on bidirectional propagation alone. The algorithm sometimes defined a bundle threshold lower than the value obtained by manual analysis. In 80 out of 100 such cases examined, subsequent inspection (by viewing the same movies of the recorded activity, but plotted on a more sensitive scale) revealed the presence of an apparent bundle signal not initially detected with manual analysis. For each of the five preparations included in the comparison (preparations 2-6 in Fig. 4B), the thresholds identified by the algorithm were: 0.93 ± 0.34, 1.46 ± 0.52, 1.39 ± 0.50, 0.86 ± 0.31, and 1.17 ± 0.44 µA (mean ± SD across electrodes for each preparation). Note that thresholds manually identified by different observers also differed somewhat (Fig. 5C). On a subset of the data reported in Fig. 5A, the correlation between bundle thresholds defined by two observers was 0.97, a value comparable to the correlation observed between the automated and manual methods.

**Figure 5:**
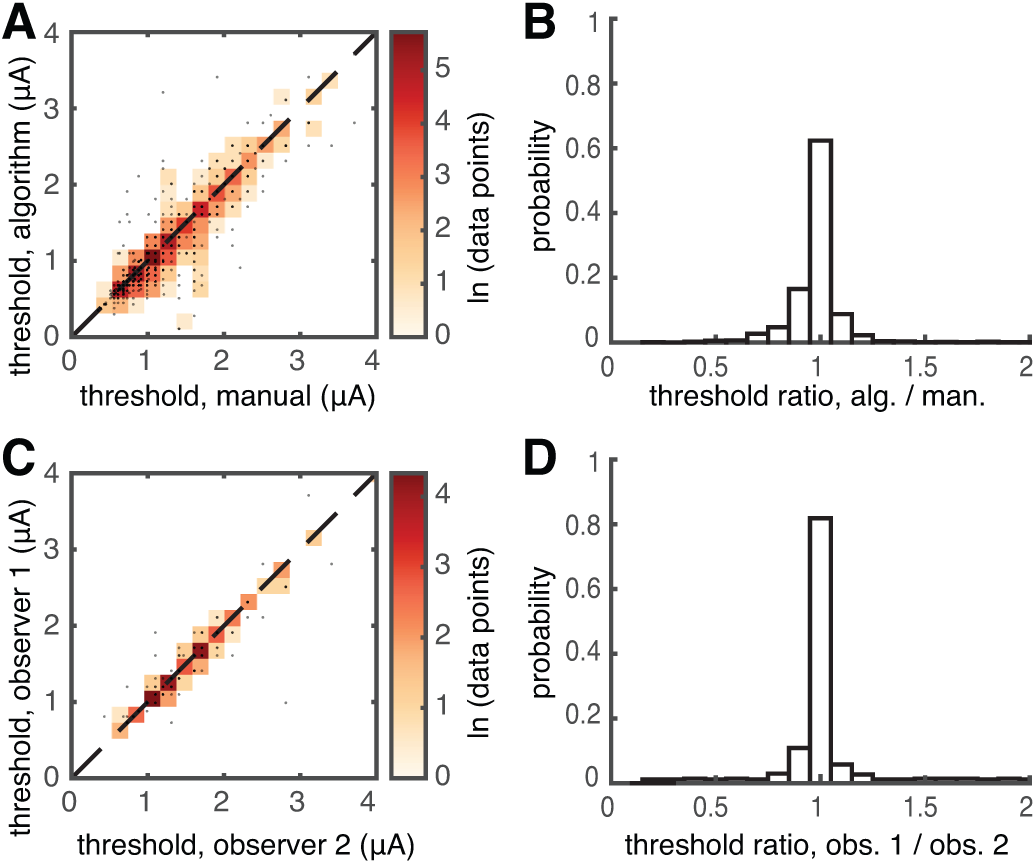
Automated and manual axon bundle threshold estimation yield similar results. A. Scatter plot compares the bundle activation thresholds identified by the algorithm and manual inspection. Color is proportional to the logarithm of the density of points. The correlation between the two methods was 0.93 for the 1885 tested electrodes shown (5 preparations). B. Histogram shows the ratio of the thresholds identified by the algorithm and the manual observer, revealing that 88% of the estimates obtained from the two approaches were within ±10% (3 central bins). C. Scatter plot shows variability in manual inspection. The correlation of threshold values reported by two observers was 0.97. D. Histogram shows the ratio of the thresholds identified by the two observers, revealing that 95% of estimated threshold values were within ±10%.

Automated analysis identified bundle thresholds with high sensitivity. Sensitivity was assessed by measuring the ability of the algorithm to detect signals from single axons in synthetic data. Synthetic data was constructed from electrical images of individual RGCs, modified to produce bi-directional propagation, and added to voltage recordings obtained with a stimulus amplitude immediately below the identified bundle threshold (see Appendix). The synthetic data consisted of 154 electrical images added to post-stimulation recordings from two preparations. The 2 individual RGC signal was either added to 100% of trials threshold, manual (μA) (Fig. 6A) or 50% of trials (Fig. 6B). In 78% of cases with signals added to 100% of trials, the algorithm identified this synthetic single-axon signal correctly (Fig. 6A). In 18% of cases, the added signal caused the algorithm to underestimate the actual bundle threshold, and in 3% of cases, the algorithm did not identify the added signal. In a few examples (2%), the added signal caused the algorithm to identify the bundle at a higher threshold than it did without the added synthetic signal. For the synthetic data with single-axon signals added to 50% of the trials, the algorithm identified the added signal correctly in 62% of cases, and missed it in 20% of cases. In 16% of cases, the algorithm underestimated the bundle threshold. These results suggest that bundle thresholds identified by the algorithm were accurate in a large majority of cases, and if anything erred in the direction of underestimating the bundle threshold. Thus, the remaining analyses (except for Fig. 9) were carried out using the algorithm to detect the bundle threshold.

**Figure 6:**
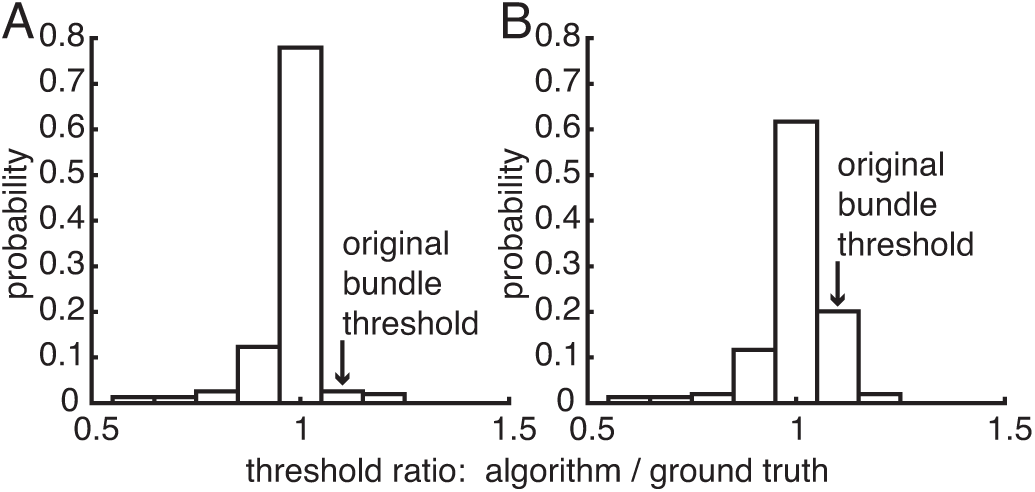
Sensitivity of automated bundle detection to individual axon signals in synthetic data. A. When the synthetic single axon signal (see Methods) was added to 100% of trials one amplitude step lower than the originally estimated bundle threshold, it was detected correctly in 78% of cases tested. B. When the single axon signal was added to 50% of trials, it was detected correctly in 62% of cases tested.

### Somatic activation is sometimes possible without axon bundle activation

Electrical stimulation at current amplitudes near bundle activation threshold often permitted the selective activation of individual RGCs. Activation of RGCs was determined by examining the voltage recordings on the stimulating electrode and surrounding 6 electrodes, immediately following stimulation, for the presence of large-amplitude spikes with a spatio-temporal waveform closely matching spikes from an individual RGC identified separately during visual stimulation (see Methods). The eliciting of these easily-identified spikes by electrical stimulation will be referred to as somatic activation, although the actual site of activation may well be the axon initial segment (Sekirnjak et al., 2008; Fried et al., 2009) or elsewhere. In each preparation, somatic activation thresholds for RGCs were in approximately the same current range as bundle activation thresholds (Fig. 7).

**Figure 7:**
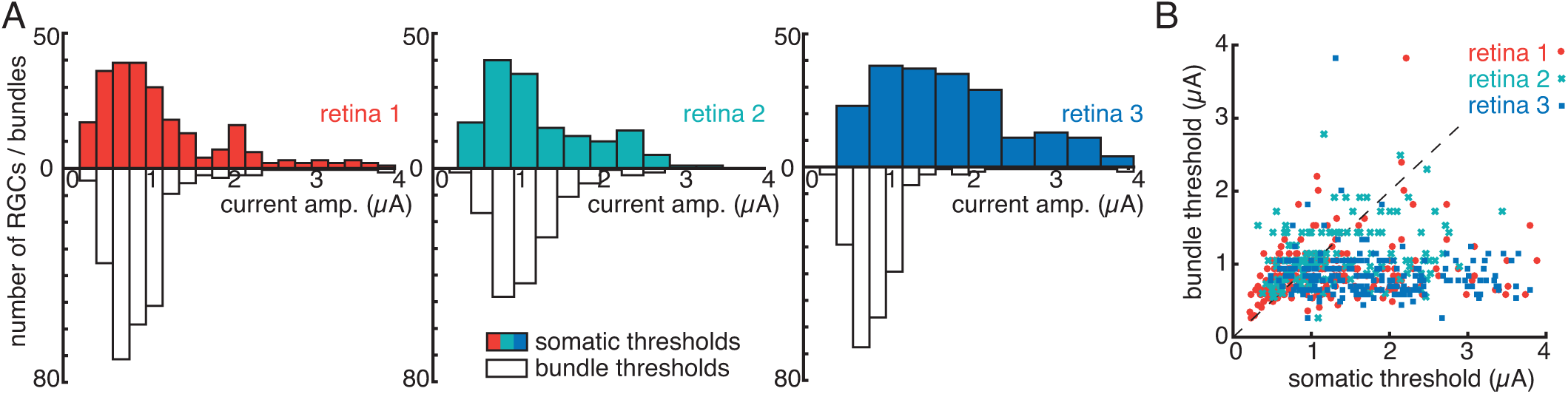
RGC and bundle threshold comparisons for three retinal preparations. A. Colored histograms show RGC activation thresholds for three preparations with eccentricities of 48.2°, 58.1° and 58.1°. White histograms show axon bundle thresholds for the electrodes used to stimulate the same RGCs. B. Scatter plot shows the RGC somatic threshold vs. bundle threshold at that electrode. In aggregate, 45% of all electrodes that were able to activate a single RGC (i.e. 17% of all electrodes on the array) were able to do so without bundle activation.

Given the similar activation thresholds for axon bundles and RGC somas, the fundamental problem for development of high resolution epiretinal prostheses is whether individual RGCs can be activated at their somas without activating bundles. This was tested by evaluating the fraction of electrodes on which such activation was possible. On average across three preparations at temporal eccentricities of 48.2°, 58.1° and 58.1°, 97% of electrodes were able to elicit some electrical activity, as revealed by bundle activation (see Fig. 4). Analysis was restricted to the 243 (47%), 150 (29%) and 199 (39%) of electrodes on which somatic activation of a single RGC was detected on the stimulating electrode and its neighbors (see above). Of these electrodes, 115 (47%), 73 (49%) and 79 (40%) were able to activate a single RGC using a stimulation current level at or below bundle activation threshold for that electrode. Thus, in aggregate, 45% of all electrodes that were able to activate a single RGC (i.e. 17% of all electrodes on the array) were able to do so without activating bundles.

The impact of these results on the ability of a prosthesis to reproduce the neural code of the retina were further examined by analyzing the proportion of RGCs of each type that were activated selectively at or below bundle threshold. As an example, consider the activation of ON parasol RGCs from one preparation (temporal eccentricity 48.2°). In this preparation, light responses were obtained from 116 ON parasol cells. Based on the mosaic structure of RGC receptive fields, this represented 97% of the total number of ON parasol cells presumably overlying the electrode array, assuming a healthy retina. Among these cells, 78 (67%) were activated selectively, i.e. without other detectable somatic activation. The RFs of the selectively activated cells, which represent their normal encoding of visual space, densely sampled the area covered by the electrode array (Fig. 8A, blue). Among these cells, 37 (47%) exhibited somatic activation thresholds less than or equal to bundle activation threshold on one or more electrodes that stimulated it most effectively. Correspondingly, the RFs of these cells sampled the area of retina over the electrode array more sparsely (Fig. 8B, blue). The fraction of recorded cells that could be activated without activating bundles varied widely across RGC types and preparations (Table 1) (see Discussion). In summary, the collection of RGCs that could be activated selectively with stimulation currents below bundle threshold formed a patchy representation of the visual scene.

**Figure 8:**
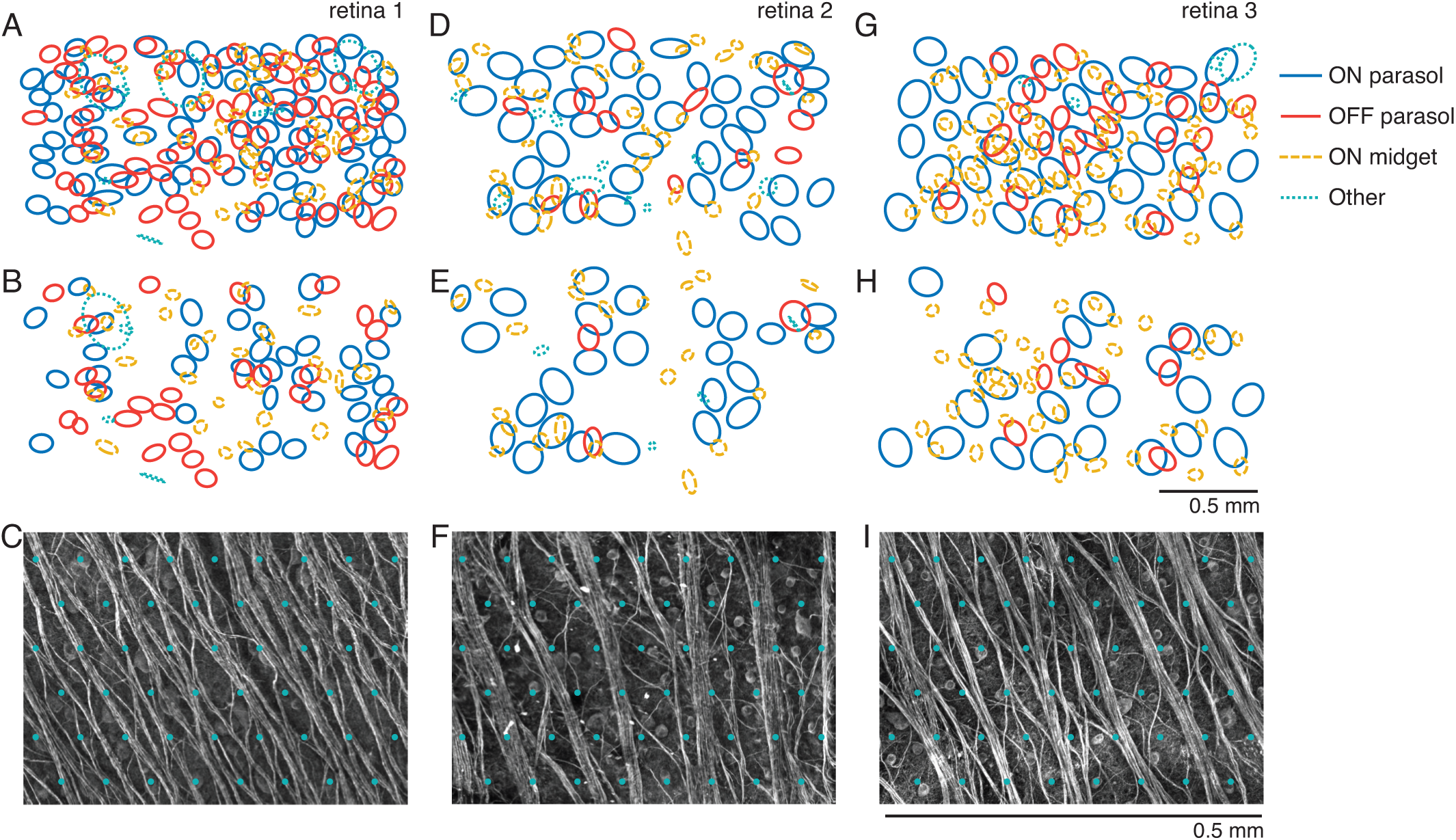
Visual receptive fields of RGCs that can be activated with and without bundle activation. The columns (A-C, D-F, and G-I) correspond to the retinal preparations described in the text with eccentricities of 48.2°, 58.1° and 58.1°, respectively. Top row (A,D,G) shows the receptive fields of the cells that can be activated at their somas without activating other nearby somas. Receptive fields are separated into ON and OFF parasol cells, ON midget cells, and other cells, which includes OFF midget cells, small bistratified cells, and cells for which the anatomical identity is unknown. Middle row (B,E,H) shows the receptive fields of the cells that can be activated without activating bundles. Bottom row (C,F,I) shows zoomed images of axon bundles in each preparation, with respect to a grid of electrodes (green overlay, arbitrary alignment) with spacing equal to that used in the experiments.

### Central recording

The applicability of these results for a clinical prosthesis in the central retina was evaluated by recording and stimulating in the Raphe region, which occupies 0-20° eccentricity on the temporal horizontal meridian. The Raphe is avoided by peripheral axon bundles that arc around the fovea as they head toward the optic disc, and thus exhibits low axon bundle density compared to other central locations (Fig. 9A). Following the analysis above, in one Raphe recording (7.3° eccentricity), 421 (98%) of 430 analyzable electrodes produced some degree of electrical activation, as assessed by bundle activity. Selective somatic activation of individual RGCs was observed with 93 (22%) of the electrodes. Among these, 70 (75%) produced selective somatic RGC activation at or below bundle threshold (Fig. 9C), compared with 45% of electrodes that did so in the peripheral recordings (Fig. 7B). Thus, in the Raphe, 75% of all electrodes that were able to activate a single RGC (i.e. 16% of all electrodes on the array) were able to do so without bundle activation. Separate analysis was not performed on different RGC types (e.g., Table 1), because the light response data collected were insufficient for reliable cell type classification.

**Figure 9:**
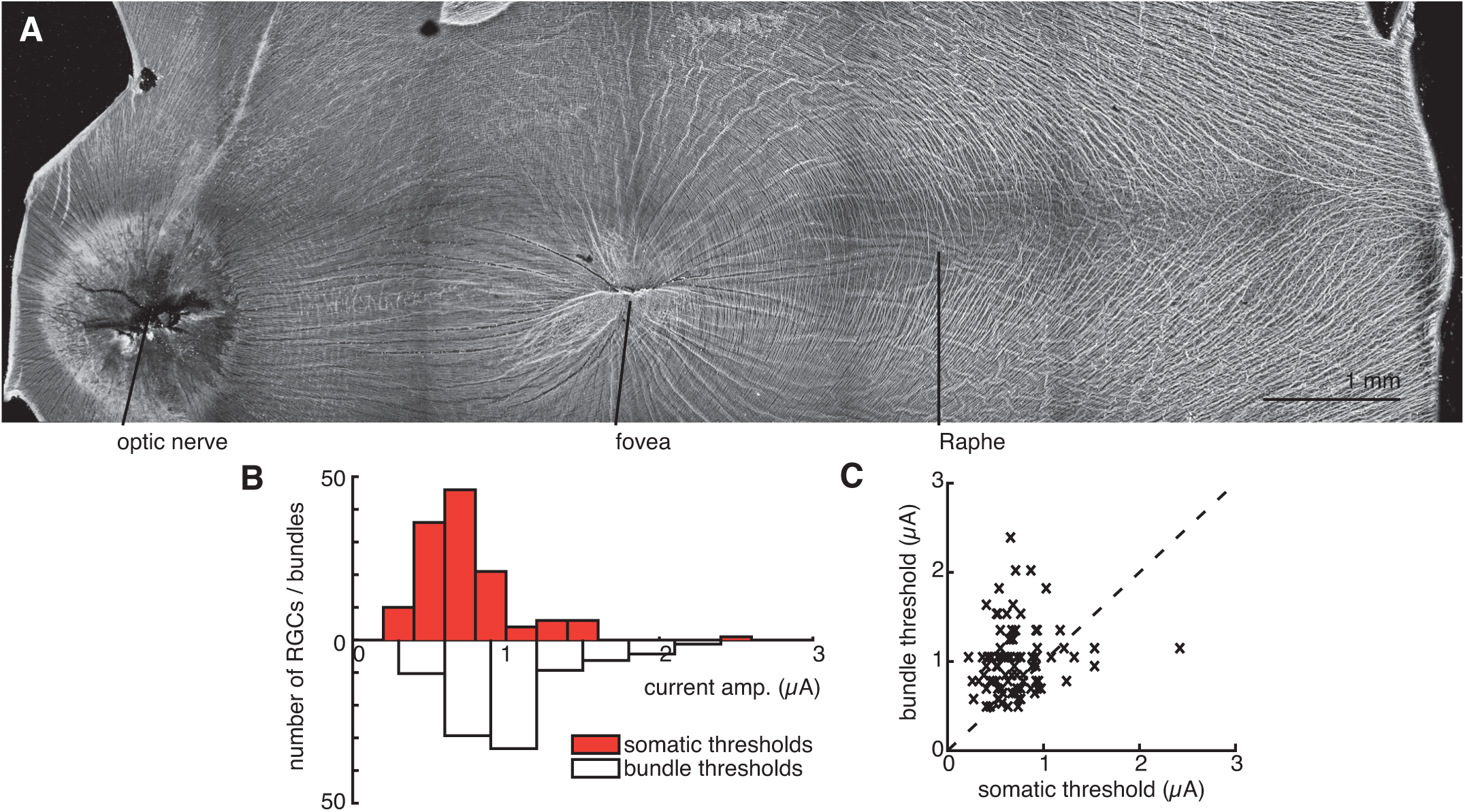
Stimulation in the Raphe region. A. Immunolabeling reveals the Raphe region of the primate retina, with a relatively low density of axons. B-C. Scatter plot and histogram show the RGC somatic threshold vs. bundle thresholds at the same electrodes (see Fig. 7). 75% of all electrodes that were able to activate a single RGC (i.e. 16% of all electrodes on the array) did so without bundle activation.

## 4 Discussion

The present findings reveal that although axon bundle activation occurs at stimulation current levels similar to those that produce somatic activation, a minority of RGCs can nonetheless be activated selectively with epiretinal stimulation, without the unwanted activation of bundles. Thus, it may be possible to precisely reproduce a portion of the neural code in an epiretinal prosthesis (Jepson et al., 2014a), while avoiding the distortions produced by axon activation in current clinical devices (Nanduri, 2011; Nanduri et al., 2012). Below, some of the implications for the development of future prostheses, and caveats of the present findings, are explored further.

### Application

To apply these findings in a clinical prosthesis, and accurately reproduce the neural code in a subset of RGCs, would require a system architecture quite unlike that of present-day devices. First, it would be necessary to calibrate the stimulation current levels of the device to target the subset of RGCs that can be selectively activated. This would presumably involve incorporating circuitry into the device to record the activity elicited by electrical stimulation, processing the recordings in a manner similar to the analysis presented here, and then adjusting stimulation current levels to the optimal values for selective RGC activation. Second, reproducing the neural code of the retina would require identifying distinct RGC types recorded, and using models of visual response to determine the appropriate firing pattern of each RGC based on its type and location with respect to the input image (Pillow et al., 2008). This would also require developing methods to identify RGC types using only intrinsic properties of cells (Richard et al., 2015), because light responses would not be available for this purpose in the blind patient. Third, spatially patterned stimulation (Jepson et al., 2014b) or optimized stimulus waveforms could potentially be used to optimize the selectivity of stimulation. Finally, this precisely calibrated stimulation approach would likely need to be adjusted periodically over time, as the implant shifts relative to the tissue or if an immune response occurs near the implant (Polikov et al., 2005). This vision of a future device raises many technical issues, including close apposition of the implant to prevent current spread, stability over time, and size and power dissipation constraints associated with high-bandwidth electrical recording and mimicking of natural retinal responses. Although no neural interfaces with such capacities have yet been developed, the relatively accessible retina and its well-understood function may provide the ideal setting for a first attempt to interface to nervous system circuitry at its native resolution.

### Central retina

In much of the central retina, the thick layer of axons overlying RGC somas would almost certainly make selective activation of RGCs without activating axon bundles more difficult than in the peripheral recordings presented (Fig. 7–8). Given these factors, the Raphe region of the central retina (Fig. 9), with its low axon density, may represent the ideal target location for a high-resolution epiretinal prosthesis. The results of recording in the Raphe support this possibility (Fig. 9), demonstrating that the fraction of electrodes that could produce selectively activated RGCs was comparable to or higher than the fraction observed in the peripheral retina. Recent emphasis with clinical devices has been on targeting the fovea, rather than the Raphe or other more peripheral areas, because of superior clinical outcomes (Ahuja et al., 2011; Humayun et al., 2012; Stingl et al., 2013a). However, the comparisons in the clinic have been performed using devices that activate RGCs indiscriminately and simultaneously over large areas. This kind of activation produces a visual signal that is never observed in nature: in the healthy retina, RGCs of 20 types signal information in their precise and distinct spatiotemporal firing patterns (Field et al., 2007). The present results suggest an entirely different approach to epiretinal electrical stimulation: reproducing the neural code cell-by-cell and spike-by-spike, in a subset of RGCs, in a central but perhaps not foveal region of the retina. Thus, the unanswered clinical question is: would a partial but precise artificial visual image in the Raphe region provide clinical outcomes superior or inferior to unnatural, indiscriminate stimulation of RGCs in the fovea? Answering this question will probably require development of the technology and testing in vivo, although computational modeling currently underway may provide some clues.

### Imaging

A possible alternative approach to avoiding axon bundle activation would be imaging of bundles before device implantation, using optical coherence tomography (Kocaoglu et al., 2011; Shemonski et al., 2015). This could potentially yield a simpler or faster way to identify bundle locations and the electrodes that would most likely activate them. In the present data, images of bundles (e.g., Fig. 2A, 8C, 8F, 8I) obtained with immunohistochemistry after recording were compared with bundle activation thresholds (e.g., Fig. 4). The correlations between image intensity summed over a 15 µm radius of each electrode and bundle threshold at that electrode were weak (0.34) but statistically significant for one low bundle density preparation (e.g., Fig. 2A), and near zero for preparations with higher bundle density (e.g., Fig. 8C). Thus, although bundle position estimated from imaging seems to influence bundle activation thresholds in some cases, other factors such as thickness of the inner limiting membrane or proximity of the tissue to the electrode array are also important. Given that bundle activation thresholds and somatic activation thresholds are similar, these observations suggest that bundle avoidance will be much more effective with electrical calibration of the kind performed here than with imaging.

### Spatial resolution

The present results were obtained using stimulation through individual electrodes on the multi-electrode array. However, spatial manipulations could improve the selective recruitment of cells over axons. In fact, multi-electrode patterned stimulation has been shown to allow selective activation of a target cell without activating nearby cells, when single electrode stimulation failed (Jepson et al., 2014b), using the same experimental methods. In addition, the 60 µm pitch of the electrode array limited the resolution with which stimulation locations could be probed. It is possible that higher selectivity could be obtained with denser arrays (see (Radivojevic et al., 2016)). These considerations suggest that the present results represent a conservative estimate of how effectively single RGCs can be activated without evoking bundle activity.

### Temporal modulation

Previous results (Freeman et al., 2010; Weitz et al., 2013; Boinagrov et al., 2014; Weitz et al., 2015) have shown that changes in the waveform of electrical stimulation can target bipolar cells over RGC axons, in a manner that substantially affects the recruitment of axon activity, due to the different integration times of different cells and compartments. Although the long pulses that elicit network-mediated responses are not appropriate for fine-grained reproduction of the temporal structure of retinal spike trains, the general concept of improved temporal stimulation patterns remains open to further exploration, and again suggests that the current findings represent a conservative estimate of how selectively activity can be evoked in RGCs.

### Completeness

Ultimately, understanding the full artificial signal created by electrical stimulation, including the complete set of cells that can be activated without producing unwanted bundle activity, would require recording reliably from every cell and axon in the vicinity of the electrode array. This goal represents a substantial challenge for current laboratory recording technology, let alone implantable hardware. Calcium imaging approaches increasingly provide an attractive technology option (Weitz et al., 2015) in the laboratory, though at present they are limited to detecting bursts of spikes with low temporal precision. The techniques used here probably represent the closest approximation to recording complete spike trains from complete neural populations in the mammalian nervous system (Frechette et al., 2004; Field et al., 2010; Segev et al., 2004), and in principle could be performed using the electrodes a prosthesis. However, significant uncertainties remain. Based on the known mosaic organization of the major RGC types in primate retina (Dacey, 2004; Chichilnisky and Kalmar, 2002; Frechette et al., 2005), a substantial fraction of cells overlying the electrode array were not recorded, to a degree that depended strongly on cell type (Table 1). It is possible that activation of these cells or their axons was not detected by the present methods. Indeed, previous work using the same methods but smaller electrode arrays likely underestimated the influence of axons by analyzing only somatic spikes without access to propagating signals over a large array (Jepson et al., 2014a). In the present work, control analysis with synthetic data (see Results) revealed that activation of single axons was usually detectable (this would likely not be the case with calcium imaging (Weitz et al., 2015)). However, the axon signals used for synthetic data were obtained from the cells that were identified during recordings with visual stimulation; it is possible that axons with smaller recorded signals were activated by electrical stimulation. Furthermore, it is possible that selective somatic activation of a single RGC would be over-estimated by the present methods, if other RGCs produced somatic spikes too small to detect. On the other hand, it is possible that RGCs with small spikes were selectively activated at current levels below bundle threshold, but could not be identified with confidence using the present methods, particularly because of the large electrical stimulation artifact. It is difficult to assess the collective impact of these factors, which could bias the estimated selectivity upward or downward. In principle, if the factors that limit the efficiency of recording similarly limit the efficiency of activation (e.g., electrode and local tissue properties), then the present results on the fraction of electrodes and cells that can yield selective activation may represent a reasonable approximation to the in vivo situation relevant for a clinical device.

## 5 Appendix

### Automated axon bundle threshold detection algorithm

Three criteria were developed for detection of bundle activity, based on the spatial structure of signal propagation and the growth of the signal with stimulus current amplitude. Because axon bundles form tracks that cross the electrode array, the algorithm was designed around path discovery techniques on graphs to leverage this prior information.

### 1. Graph creation

The spatial pattern of responses to electrical stimulation was represented as a directed graph, with vertices corresponding to the electrodes, and edge weights set according to the amplitude of signals recorded on all electrodes following stimulation on each electrode. Specifically, the weight *W_ij_* of the edge joining the vertex representing stimulating electrode *i* and the vertex representing a second electrode *j* was given by:

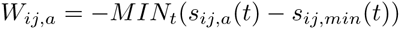

where *s_ij,a_*(*t*) and *s_ij,min_*(*t*) represent the voltage waveforms over time *t* recorded on electrode *j* (average across n = 25 trials), using stimulus current amplitude *a* and the minimum amplitude respectively on electrode *i*. *M IN_t_* represents the minimum of the voltage recorded over 5 ms after the application of the stimulus pulse. The minimum was used because the largest magnitude voltage deflection observed in axon bundle signals was negative. The weight obtained with the maximum current amplitude will be referred to simply as *W_ij_*. These weights form an adjacency matrix *W* for the entire electrode array.

### 2. Estimated bundle paths via graph traversal

The next step estimated the position of the axon bundle(s) activated by each stimulating electrode, *i*. The start and end points for the path of the bundle were defined as the two array border electrodes with highest values of *W_ij_*, indicating the exit point of the traveling bundle signal from the array area. The start and end points could be on any of the four array borders, but not on the same border as one another (Fig. 10A). For each stimulation current *a*, A* pathfinding (Hart et al., 1968) was applied to define a path between the start and end points, minimizing the sum of a cost function and heuristic function. The cost of traveling from vertex *k* to vertex *j* was defined as *W_kj,a_ − d*(*k, j*) where *d*(*k, j*) is the physical distance between electrodes *k* and *j*. The heuristic function value for vertex *k* was the minimum number of nearest neighbor electrode steps required for a path joining electrode *k* to the end point. The result of the A* algorithm was a path *P_i_, a* associated with stimulating electrode *i* and stimulation current amplitude *a*, connecting the start and end points. The path obtained with the maximum stimulus current amplitude will be referred to simply as *P_i_*.

**Figure 10:**
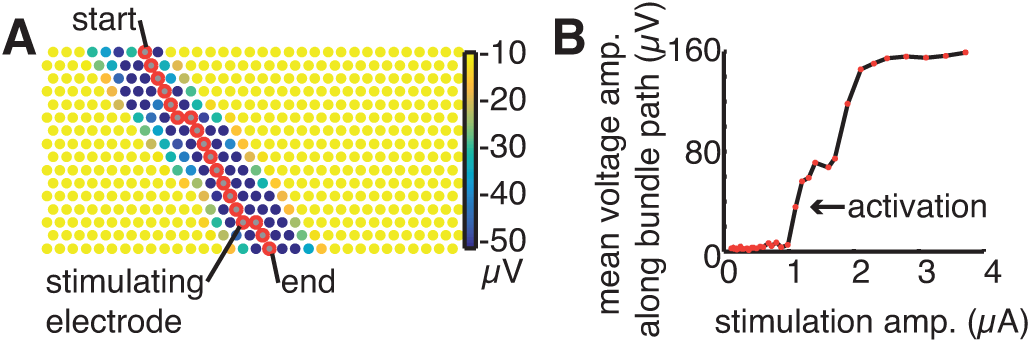
Bundle paths and signal growth identified with graph traversal. A. Electrodes on the array are represented by circles, colored by the minimum recorded voltage on each electrode during the 5 ms following electrical stimulation. Start and end points were chosen as the electrodes on the array borders with the largest recorded signal, and a path (red) was identified joining these electrodes. B. Mean voltage deflections recorded on electrodes in the bundle path (A) as a function of stimulation amplitude were used to identify the bundle activation threshold. Bundle activation was identified when the squared difference of sequential points on the curve exceeded a fixed threshold. Data from same preparation as Fig. 3.

### 3. Growth criterion

At each stimulating electrode, the growth in the recorded voltage along an estimated bundle path was examined as a test for bundle activation. Specifically, the mean of the weights *W*_*ij,a*_ along the electrodes *j* in the path *P_i_, a* was examined as a function of stimulation amplitude *a* (Fig. 10B). If the squared difference between mean voltages obtained at two consecutive stimulation levels exceeded a fixed threshold, the higher of the two stimulus levels was identified as producing bundle activation according to the growth criterion (see below). The squared difference threshold was set to 60 µV^2^, the value that produced the best match to human estimates in 5 data sets.

### 4. Graph partitioning

The electrode array was then partitioned into disjoint spatial regions reflecting axon bundle geometry, for analysis of signal propagation. First, entries in the adjacency matrix *W* that were not connected by a path *P_i_* (Step 2), were zeroed (Fig. 11B). Next, a symmetrized adjacency matrix *W_u_* = *WW*^*T*^ + *W*^*T*^*W* was calculated (Fig. 11B) (Satuluri and Parthasarathy, 2011; Malliaros and Vazirgiannis, 2013). The graph was then partitioned into disjoint groups of electrodes using a spectral decomposition of the modularity matrix (Newman, 2006). An additional constraint was imposed such that each group of electrodes was contiguous in space (connected by physically neighboring electrodes), and intersected at least two borders of the electrode array (Fig. 11C-D). These constraints were imposed by merging disconnected groups of electrodes that were closest in modularity until the preceding conditions were satisfied.

**Figure 11:**
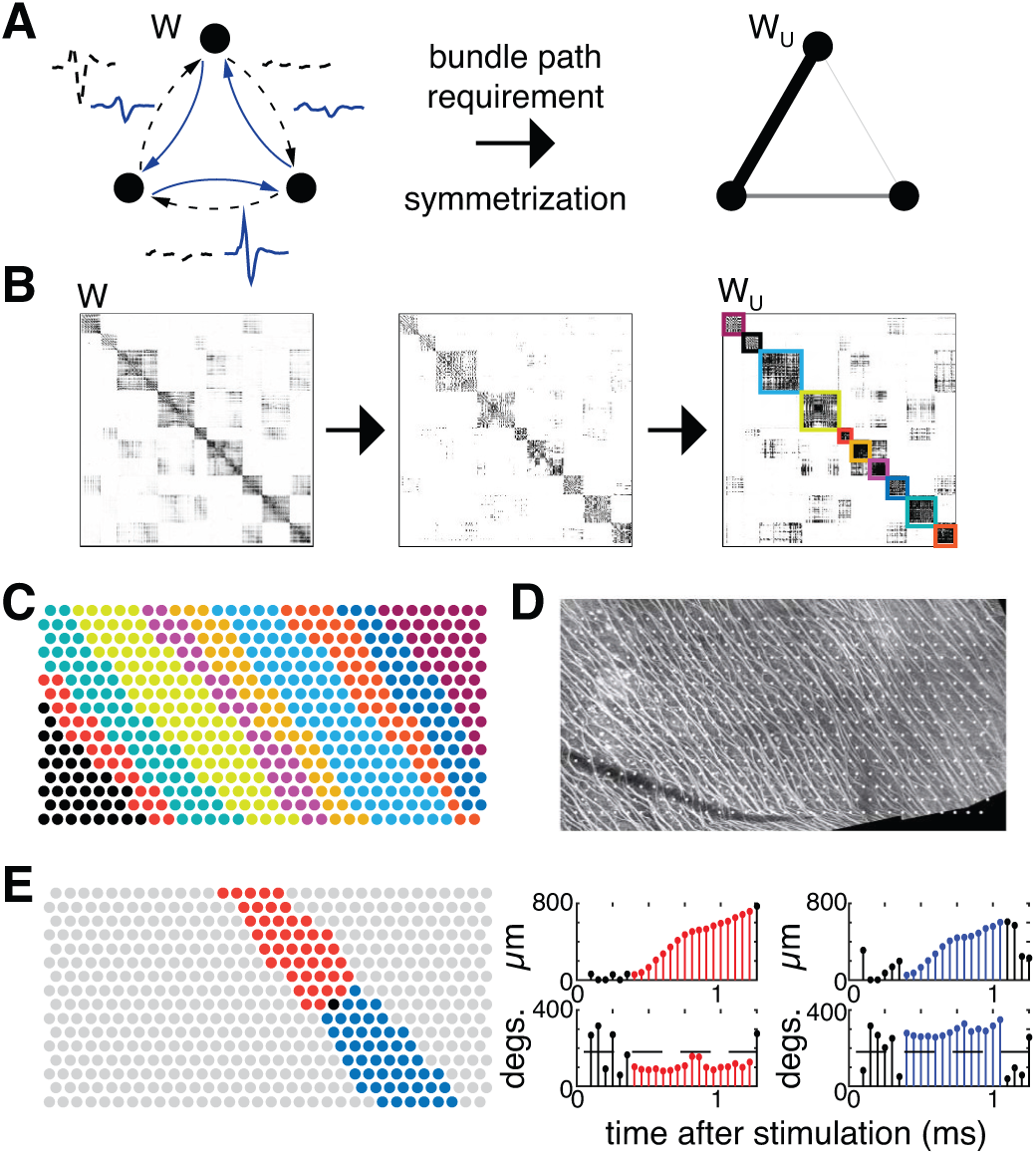
Illustration of graph creation, partitioning, and bidirectional propagation testing to determine bundle activation thresholds. A. Left: mean waveforms recorded on all electrodes after stimulation at each electrode were used to construct a complete, directed, weighted graph. The graph is transformed by zeroing all edges not connected by a path produced by electrical stimulation, then symmetrized. Right: Weights in the resulting undirected graph are represented by the thickness of connecting lines (thicker line corresponds to a stronger connection). B. Left: Initial 512x512 adjacency matrix for one electrical stimulation data set. Middle: Entries in the preceding adjacency matrix not connected by a path were zeroed. Right: The adjacency matrix was symmetrized and the graph partitioned on the basis of its modularity. Groups of electrodes that were identified are outlined using distinct colors. C. Groups of electrodes are revealed using the same colors as B. D. IHC image of axon bundles for same preparation as C, with electrodes overlaid. The groups of electrodes from C conform to the geometry and direction of labeled axon bundles. E. Visualization of bidirectional propagation testing. Left: the group of one particular stimulating electrode (in black), is bisected, resulting in two bands (red, blue). Right: stem-plots show displacement and angle of movement of the center of mass of elicited activity in each of the two bands. Consecutive time samples with bidirectional propagation are highlighted in red and blue, respectively.

### 5. Bidirectional propagation

At each stimulating electrode, a test for bidirectional propagation of the electrical response signal was performed, using the electrode partition to identify diverging movement of the pattern of activity from the location of the stimulating electrode. For each stimulating electrode, the corresponding group of electrodes (defined in Step 4) was bisected about the line through the stimulating electrode that was perpendicular to the least-squares fit line through the electrode positions in the group. This yielded two bands of electrodes running in opposite directions from the stimulating electrode (11E, red and blue bands). When an electrode was exactly on the border between two groups, the union of the bordering groups was taken before splitting into bands. Then, at each stimulation amplitude, signal propagation was tracked in space and time for each of the two bands separately. First, all but the five most negative samples in the mean recorded waveform on each electrode over 5 ms following stimulation were zeroed. Second, at each time sample, a spatial center of mass (Li et al., 2015) was computed for each band of electrodes, using the partially zeroed waveforms for weights. Third, sequences of consecutive time points were identified for which the distance from the center of mass to the stimulating electrode monotonically increased (sequences are colored red and blue in Fig. 11E), while the angle of the movement between consecutive time points remained within 90 degrees of the least-squares regression line through the electrode positions of the group (Fig. 11E, right). This identified sequences of consecutive time points during which the center of mass of activity moved away from the stimulating electrode approximately in the direction of the band. All such time sequences were then examined further to probe the extent of the propagation in space. If the net movement of the centers of mass exceeded half the distance from the stimulating electrode to the array border in both directions, within any of the sequences of consecutive time points identified above, the stimulus was identified as producing bundle activation according to the bidirectional propagation criterion. This criterion was selected to most closely mimic human estimates of bundle activation. If propagation in the direction opposing the normal direction of spike propagation (one of the two bands) exceeded half the distance from the stimulating electrode to the array border, the stimulus was identified as producing bundle activation according to the back-propagation criterion.

### 6. Assignment of bundle threshold

Bundle activation criteria were then applied as follows:

1: If the stimulating electrode was within two electrodes of the array border, the growth criterion alone was used. 2: For other stimulating electrodes, if the back-propagation criterion indicated bundle activation at stimulus amplitudes lower than the growth criterion, the former was used. 3: In all other situations, the growth criterion and bi-directional propagation criterion were tested, and the criterion indicating activation at a lower stimulus current level was used.

Bundle activation threshold was defined as the midpoint between the lowest current level at which bundle activation was detected and the next lower current level tested.

### Synthetic data for evaluation of the sensitivity of the algorithm

Synthetic data was created to test the sensitivity of the axon bundle detection algorithm. Signals from individual RGCs were added into mean-subtracted data recorded following electrical stimulation. The synthetic data were built to test if the algorithm was sensitive to the presence of a single activated axon under normal noise conditions. First, an electrical image obtained during visual stimulation was modified such that the spike initiated at an arbitrary electrode along its axon, chosen to be the “stimulating” electrode. The minimum of the waveform at the stimulating electrode defined the start time (*t*_0_), and waveforms on all other electrodes with minima at times < *t*_0_ were reversed in time. The modified electrical image was then added to either 100% or 50% of trials, with a latency of 250 µs following the stimulus, which is typical for RGC activation. Since the mean of all trials was used to detect a bundle, the data with EIs added to only 50% of trials contained signals that were smaller than typically recorded for a single RGC. The modified electrical image was added at the next stimulation amplitude lower than the amplitude at which bundle activation was previously detected. The range of mean voltages of the added EI signal used to construct the synthetic data had a similar distribution as the EI signal obtained from all recorded cells.

## 6 Supplemental Material

Supplemental data show the recorded voltage signals as a movie overlaid with the tubulin image. The propagating signal is coarsely aligned with the direction of axon bundles, and can be seen propagating unidirectionally (Supp. video 1, stimulation amplitude is 0.99 µA) or bidirectionally (Supp. video 2, stimulation amplitude is 1.91 µA), depending on the stimulation current amplitude. Data shown is the same as in Fig. 2.

## 7 Author contributions

L.E.G. and E.J.C. conceived and designed the experiments, L.E.G., G.A.G., S.Ma., and P.L. collected the data, L.E.G., K.G., S.Ma., N.B., and V.F. analyzed the data, K.G and N.B. developed the algorithm, P.H., W.D., A.S., and A.M.L. developed and supported the stimulation and recording hardware and software, L.E.G. and E.J.C. wrote the paper, all authors edited the paper, S.Mi. and E.J.C. supervised the project.

## 8 Acknowledgements

Research reported in this publication was supported by the National Eye Institute of the National Institutes of Health, award numbers F32EY025120 (LEG), EY021271 (EJC), and 5R21EB004410 (AML), National Science Foundation grant PHY-0750525 (AML), Stanford Neurosciences Institute (EJC, SM), a Stanford Neurosciences Institute Interdisciplinary Scholar award (GAG), Polish National Science Centre grant DEC-2013/10/M/NZ4/00268 (PH), Pew Charitable Trusts Scholarship in the Biomedical Sciences (AS). The content is solely the responsibility of the authors and does not necessarily represent the official views of the National Institutes of Health. Confocal images were acquired using the Stanford Neuroscience Microscopy Service, supported by NIH NS069375. We thank D. Palanker for inspiration, stimulating discourse, and comments on the manuscript; F. Kellison-Linn and J. Desnoyer for assistance with analysis; W. Newsome, M. Taffe, T. Moore, J. Carmena, C. Ferrecchia, and the UC Davis Primate Center for providing access to retinas; and D. Sandel, D. Chambers, and the Salk Institute Imaging Facility for technical assistance.

